# Synaptic weight decay with selective consolidation enables fast learning without catastrophic forgetting

**DOI:** 10.1101/613265

**Authors:** Pascal Leimer, Michael Herzog, Walter Senn

## Abstract

Learning can interfere with pre-existing memories that in classical neural networks may lead to catastrophic forgetting. Different from these networks, biological synapses show an early decay of long-term potentiation, combined with a tag & capture mechanism for selective consolidation. We present a 2-component synaptic plasticity model that, by an early decay and a tag & capture mechanism, enables context-dependent fast learning without catastrophic forgetting. For reinforcement learning in multiple contexts, the fast learning combined with an early weight decay maximizes the expected reward while minimizing interferences between subsequent contexts. Fast learning, enabled by a highly plastic weight component, improves performance for a given context. Between contexts this plastic component decays to prevent interference, but selective consolidation into a stable component protects old memories. As a downside of these mechanisms, learning is hampered when consolidation is triggered prematurely by interleaving easy and difficult tasks, consistent with human psychophysical experiments.

## Introduction

Depending on the context, similar situations may require different actions. For instance, when we wait at a crosswalk in continental Europe we first look to the left, but in the UK we look to the right. After living a while at a new place, the reliably acquired behavior causes a short moment of uncertainty when crossing the road the first time back home. In general, learning to correctly act in one situation may interfere with learning in a similar situation. Retaining context information imposes a challenge to the underlying neuronal network, in particular if contexts switch quickly. On the one hand, synaptic connection strengths, which were important in a previous context, need to be protected from being overwritten. On the other hand, new associations have to be learned for similar inputs that require a different action in another context. This problem is known as the stability-plasticity dilemma, and mechanisms at the network and synapse level have been proposed to tackle it (Abraham and Robins, 2005; Fusi et al., 2005; Carpenter and Grossberg, 1988).

Catastrophic forgetting is a problem in neuronal networks, but not so much in human continual learning (French, 1999; Kumaran et al., 2016). In classical neuronal network models, the connection strengths are adapted to learn new associations in the current context regardless of the importance of the connections in previous contexts. For strong overlaps of the input patterns, ongoing learning results in memories which are forgotten on a catastrophically short time scale. To prevent catastrophic forgetting, memories can be consolidated at the level of the system or the synapses. In systems consolidation, memories are suggested to be transferred to other brain areas for long-term storage such as the hippocampus (Squire and Alvarez, 1995; McClelland et al., 1995; Roxin and Fusi, 2013). In synaptic consolidation, the fast decay of the so-called early long-term potentiation/depression (early LTP/LTD) was shown to be prevented by a synaptic tagging & capture mechanism (Frey and Morris, 1997; Sajikumar and Frey, 2004; Morris, 2006; Redondo and Morris, 2011; Shires et al., 2012; Bosch et al., 2014). Synaptic models with a cascade of internal states of progressively longer retention times were shown to prevent an exponentially fast forgetting while being continuously exposed to stimuli (Fusi et al., 2005; Benna and Fusi, 2016; Kaplanis et al., 2018). Forgetting can also be counterbalanced by reducing plasticity for weight configurations that are important in previous contexts, but assessing the importance of a synaptic weight requires additional information and memories (Kirkpatrick et al., 2017; Zenke et al., 2017; Aljundi et al., 2018).

The passive decay of the early LTP/LTD is typically seen as a weakness of the synapses that needs to be counteracted. Instead, forgetting may itself be beneficial (Brea et al., 2014). We suggest that this passive forgetting is the expression of minimizing the interference between subsequent contexts in the presence of fast learning. The fast synaptic plasticity may enable the retention of information in working memory (Mongillo and Denève, 2008) that must fade out in time to prevent cross-talks in a subsequent context where similar stimuli may need to be differently processed. The passive forgetting further allows for selectively retaining relevant information by the tag & capture consolidation (Frey and Morris, 1997). Various phenomenological models for the tag & capture mechanisms exist (Clopath et al., 2008; Barrett et al., 2009; Ziegler et al., 2015). But how can the ideas of fast learning, passive weight decay and synaptic consolidation for minimizing cross-talks be captured in a simple normative theory of synaptic plasticity and learning?

Here we suggest a 2-stage model of synaptic modifications and consolidation that maximizes the expected reward in a reinforcement learning context through stochastic gradient ascent. For simplicity and different from previous models (Fusi et al., 2005; Clopath et al., 2008; Barrett et al., 2009; Benna and Fusi, 2016; Kaplanis et al., 2018; Ziegler et al., 2015), we do not consider internal variables. Instead, our fast (‘early’) and slow (‘consolidated’) components both affect the synaptic strength. Early LTP/LTD promotes fast learning and thereby enables exploitation of reward in a reinforcement learning scenario. It necessarily includes a fast decay to prevent inferences with future contexts where similar sensory inputs require different responses. Hence, early LTP/LTD and synaptic tagging/consolidation are seen as a means to address the stability-plasticity dilemma. We suggest a normative theory of synaptic consolidation that functionally reproduces the phenomena of the multi-state models of tagging & capture (Barrett et al., 2009; Clopath et al., 2008; Ziegler et al., 2015). Our theory asserts that fast learning can be achieved in a given context with minimal interference with other contexts, provided that only strong synaptic modifications associated with peak postsynaptic depolarizations are retained. Weak synaptic modifications associated with low postsynaptic depolarizations must be extinguished on the timescale of the context duration, relating to the decay of the early LTP/LTD.

While our 2-component plasticity model boosts learning for typical context switches, it may hamper learning when the task-difficulties change too quickly. Such phenomena are in fact observed in psychophysical experiments involving stimulus or task mixing (Tartaglia et al., 2009a; Flesch et al., 2018). We postulate that the passive weight decay with the selective consolidation that allows for fast contextdependent learning is the reason why humans, unlike neural networks with simpler plasticity models, show reduced performances in these mixing experiment.

## Results

### Reward maximizing learning rule with selective consolidation

We hypothesize that early LTP and its consolidation mechanism is one of nature’s choices to deal with the stability-plasticity dilemma. To keep the benefit of fast synaptic plasticity while avoiding contextual interference, the network should only consolidate a minimum number of synapses while still be able to change enough synapses if required by novel learning. A particularly pronounced form of the stability-plasticity dilemma becomes apparent when similar tasks are learned each after the other. To illustrate this problem, we consider a learning task that extends across two subsequent contexts (Fig. 1A). The context defines the criteria according to which sensory patterns are classified, and patterns in the two contexts are similar, but not identical. They consist in written words that have to be classified in the first context according to the color of the letters, and in the second context according to the meaning of the word. Learning takes place in a single-layer network that classifies the correlated input patterns by a winner-take-all (WTA) dynamics; the first spike of an output neuron suppresses other output neurons via global inhibition (Fig. 1B).

**Fig 1.**
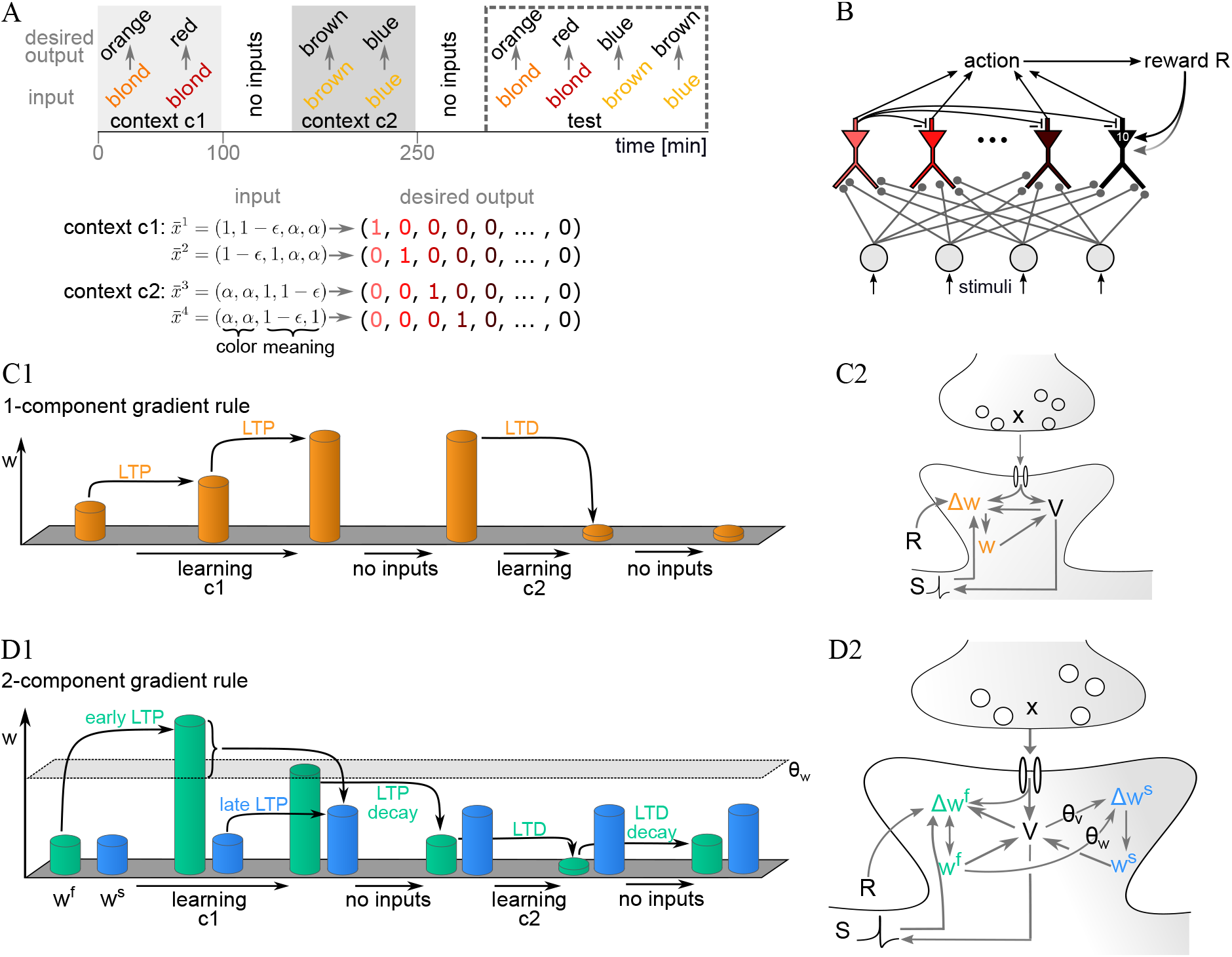
Classifying similar input patterns in different contexts fails with a 1-component plasticity model but succeeds with a 2-component model. (A) Input patterns have to be classified according to different criteria (e.g. color or letters), depending on the context. Patterns presented in one context may look similar (encoded by *ϵ*) and have strong overlaps across contexts (encoded by *α*). When being sequentially exposed to the contexts, learning in the second context interferes with the memory acquired in the first context. (B) Model network to solve the classification task using reward-based learning. The synapses from the input to the output layer are plastic (dots at line endings). Lateral inhibition in the output layer (only shown for first output neuron) enforces a winner-take-all mechanism according to which a first spike of a neuron in the output layer suppresses the possible spikes in the other output neurons right at the axonal initial segment. The winner defines an action, and a global reward signal (*R* = 1) is fed back to the network if the action is the desired one (otherwise *R* = 0). (C1) In the 1-component rule, long-term potentiation (LTP) induced in a first context c1 may be undone by long-term depression (LTD) in the second context c2, leading to forgetting of the previous weight change. (C2) The weight change Δ_*w*_ depends on the presynaptic activity *x*, the postsynaptic voltage *V*, a possible postsynaptic spike, and the reward signal *R*. (D1) In the 2-component rule, an early LTP that pushes the fast weight component *w^f^* across a threshold *θ_w_* can become consolidated in the slow weight component *w*^*s*^ upon a later strong postsynaptic activation (crossing a voltage threshold *θ_V_*, late LTP). This slow component is protected against LTD in the second context. (D2) Dependencies of the weight changes for the 2-component rule (cf. Eqs 3 and 4).

If the task is learned with a classical (1-component) reward-based learning rule, the synaptic modifications in the first context are likely to be undone during learning in the second context (Fig. 1C). To prevent this, we consider a 2-component plasticity model that consolidates appropriately selected weight changes and protects them from erasure (Fig. 1D). A fast weight component, *w^f^*, enables quick memory acquisition and acts on a short timescale, and a slow component, *w^s^*, allows for keeping a selected memory across a longer timescale. The sum of the two components determines the total synaptic strength, *w* = *w^f^* + *w^s^*.

We postulate that the dynamics of the fast weight component follows approximately the gradient of the utility function U defined by the expected reward and two penalty terms,

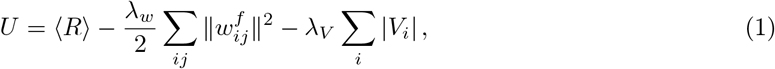

where *R* is the binary reward signal released in response to the action of the network, *V_i_* is the voltage of neuron *i*, and 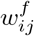 is the fast weight component of the synapse from the presynaptic neuron *j* to the postsynaptic neuron *i*. Beside the expected reward 〈*R*〉, the utility function has two penalty terms. The first term punishes high values of the fast weight components. This term results in a passive decay of the fast component such that, in the absence of new plasticity events, it converges to baseline value 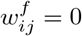. The second term punishes voltage deflections |*V_i_*| from rest at 0, and takes energy costs into account. At the same time this term helps to reduce interferences within contexts. The positive factors λ_*w*_ and λ_*V*_ represent a weighting of the two penalty terms.

To calculate the gradient of the utility function *U* with respect to 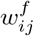 we assume that the postsynaptic neuron fires with instantaneous Poisson rate *φ*(*V_i_*), where the postsynaptic voltage is the sum of the input rates *x_j_* weighted by the effective synaptic strengths *w_ij_*,

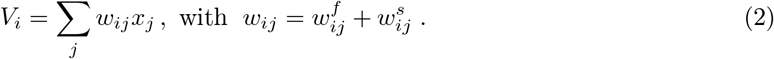

The firing rates range between 0 and *φ*_max_. Stochastic gradient ascent on the utility function *U* leads to the update of the fast weight component at repetitive time steps (assumed to be every 30 s, see Methods),

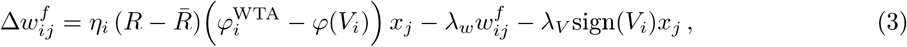

with 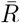 being the mean reward, and 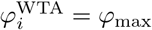 if neuron *i* is the winner neuron (*i* = *k*) and 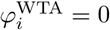 else. This time-discrete update implicitly assumes a time step Δ*t* = 1 (suppressed in Eq. 3 and below) that corresponds to the biological time of 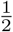 minutes. The decay term 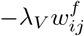, for the optimal values for context learning, causes a passive decay in the order of 1/(2λ_*w*_) = 42 biological minutes.

The activity 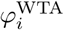 represents the target value for the instantaneous firing rate *φ*(*V_i_*). For positive reward (*R* = 1), the weights driving the winner neuron *k* are strengthened to push *φ*(*V_k_*) towards *φ*_max_, and the weights driving the non-winning neurons are weakened to push *φ*(*V_i_*) towards 0, with the opposite changes in the absence of reward (*R* = 0). The learning rate *η_i_* incorporates a positive factor necessary for the gradient property to hold (Methods). The voltage penalty causes a decrease or increase of an activated weight (*x_j_* > 0), depending on whether the postsynaptic voltage is above or below rest (*V_i_* = 0), respectively. In essence, –sign(*V_i_*)*x_j_* is an anti-Hebbian term that subtracts away the non-informative overlaps in the presynaptic activity patterns.

To store long-term memories, the fast component needs to be consolidated. We postulate that this is done when two conditions are met: (1) the instantaneous value of the fast component, |*w^f^*|, exceeds a weight threshold, and (2) the low-pass filtered postsynaptic voltage, 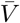, exceeds a voltage threshold. Formally, the decay 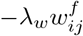 of the fast component is transferred to the slow component provided the two conditions are simultaneously satisfied,

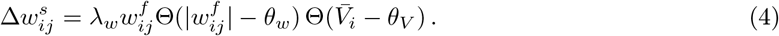

where the Heaviside step function Θ(.) is 1 if the argument is positive and 0 else.

### Preventing catastrophic forgetting with the 2-component rule

We next test the learning rule when the network performs the above mentioned learning task that is distributed across two subsequent contexts each 100 biological minutes long, interleaved by a 50 biological minutes break. To conceptualize the idea, we consider four input classes 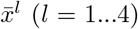, with each class requiring its desired output defined by the unique activity of a specific output neuron. Input patterns are defined as noisy samples *x^l^* around 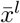 with independent Gaussian noise on the components. The mean input patterns are 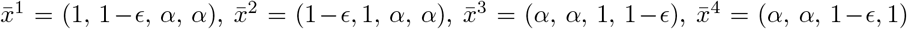, see Fig. 1A. Patterns from the first two classes are presented within the first context (c1), and patterns from the second two classes within the second context (c2). Within each context, the classes are only distinguishable by a small *ϵ* in one component, and across contexts classes have strong overlap α (close to 1). Patterns from the two classes are presented many times (200, corresponding to 100 minutes) in random order until the context switches and patterns from the other two classes are randomly presented. In response to a pattern presentation one output neuron fires first, and if this matches the desired output neuron of the class, a global reward signal (*R* = 1) is given. Otherwise reward is omitted (*R* = 0, Fig. 1B).

We first performed the learning experiment with the 1-component rule. This is the rule with only the fast component *w^f^* governed by Eq. 3 while setting the decay parameter *λ_w_* to zero and thus also *w^s^* = 0 (Fig. 1C). The other parameters were optimized for highest performance (Methods). Because the 1-component rule follows approximately the reward gradient, the associations can be learned separately within each context (Fig. 2A). However, because in the second context similar patterns are associated with different outputs, the associations learned in the first context are overwritten. The larger the overlap (*α*), the stronger the interference. When retesting the associations of the first context after learning in the second context, performance decayed roughly to a third of the original one (Fig. 2A and B, orange).

**Fig 2.**
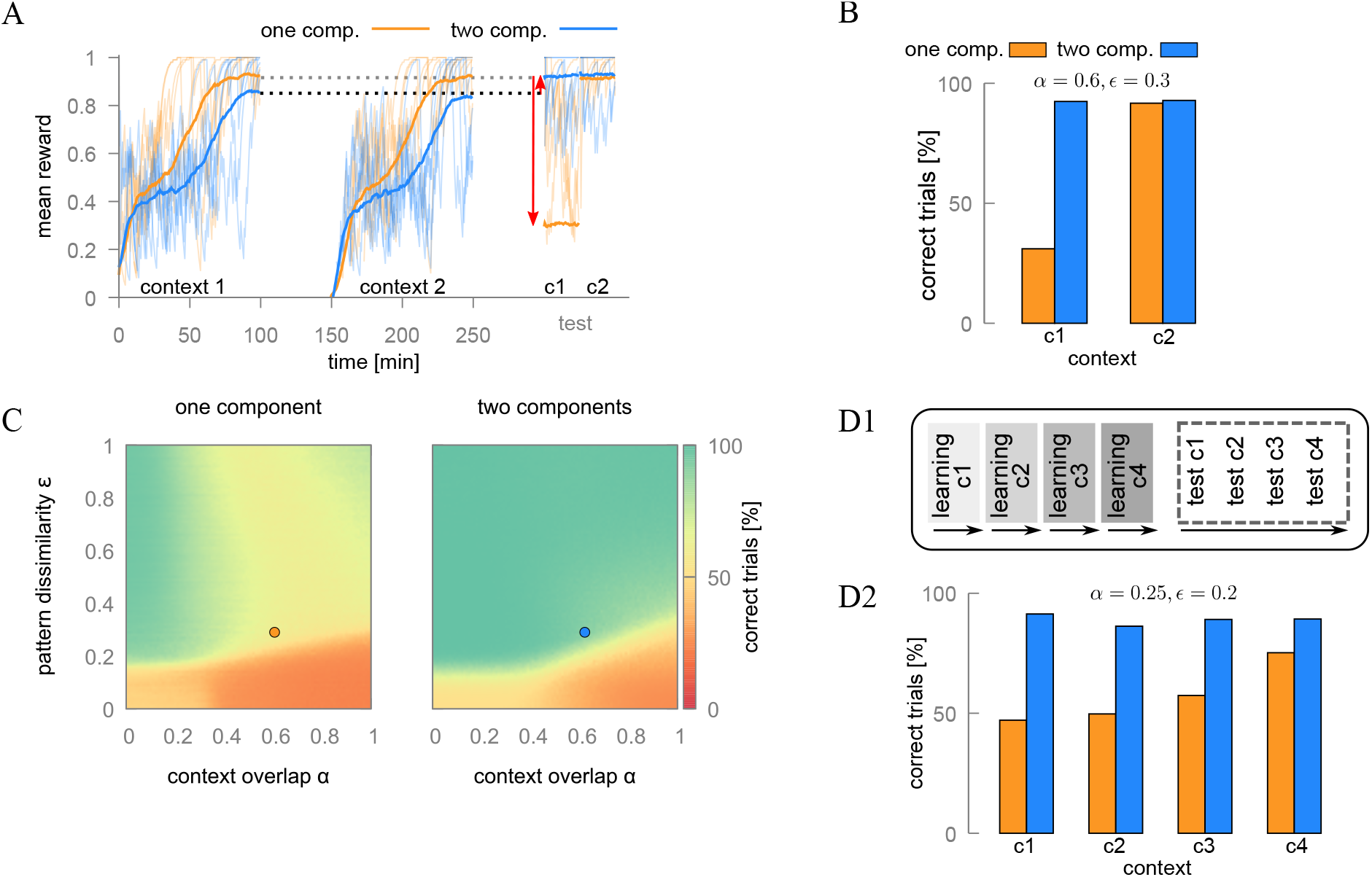
Learning similar associations in sequence causes catastrophic forgetting with a classical 1-component rule. (A) Five example traces of low-pass filtered reward 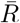 (light) and the average (bold) in two subsequent contexts, interleaved by a break without inputs. With the 1-component rule (orange) learned memories from the first context are forgotten after exposure to the second context (red downward arrow, c1), while the memories from the second context are still intact (c2). With the 2-component rule (blue) the memory performance for both contexts is improved after the second break due to the consolidation mechanisms (red upward arrow) for both contexts c1 and c2. (B) Average performance for the two contexts during the test period (c1 and c2 from panel A). (C) Average retrieval performance of all four memories for values *ϵ* and *α*. Dots show the parameters used in panel A and B. Catastrophic forgetting is mainly observed for strong context overlaps (*α* > 0.2) when using the 1-component rule. (D) When appending more contexts, each with two associations to be learned, memory performance at the final test decreases the further back context lays.

The forgetting of the previous associations can be prevented if the fast weight changes are selectively consolidated in the slow component. The informative weight changes are identified when the two conditions for the consolidation are satisfied, a large absolute value of the fast component (|*w^f^*| > *θ_w_*) and a large low-pass filtered voltage 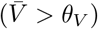. These are the filter criteria for saving the passive decay ofthe fast component (–*λ_w_w^f^*) into the non-decaying slow component (Eq. 4). Because the non-informative weight changes decay without consolidation, they do not contribute to erroneous activation of the post-synaptic neurons, and this prevents the interference-induced forgetting (Fig. 2A and B, blue).

Another benefit of the selective consolidation is that it filters out imbalanced plasticity inductions originating in the randomness of the pattern presentations and the noisy inputs. As a consequence, the slow components are less noisy than the fast components. During the delayed test period, when the fast components decayed back to baseline, we therefore obtain a better recall performance than immediately after the learning (Fig. 2A, black arrow). Hence, a break after learning improves the performance due to synaptic consolidation (that includes the decay of the last, stochastically triggered fast components), reminiscent to memory consolidation during sleep (Stickgold, 2005).

To investigate how learning depends on the pattern dissimilarity *ϵ* and context overlap *α* we repeated the experiment for many values *ϵ* ∈ [0,1] and *α* ∈ [0,1] and measured the average recall performance over all four input patterns during the test period (Fig. 2C). With the 1-component rule, already a small overlap between the patterns in the two contexts (*α* > 0.2) erased the associations learned with the first context when the second context was presented. In contrast, with the 2-component rule, the performance for the first context stayed at the original level also after exposure to the second context, even when the overlap is maximal (*α* = 1), assuming that the patterns are dissimilar enough within a single context (*ϵ* > 0.4). For more similar patterns, both learning rules show a performance decline. Extending the learning experiment by more contexts with new but similar associations supports our previous results: with 1-component rule the network gradually forgets old memories when learning new ones, but with the 2-component rule no forgetting is observed (Fig. 2D).

### Competition-agnostic synapses allow for high learning rates

Stochastic gradient rules have the intrinsic problem that the learning rate (*η*, effectively being a plasticity scaling factor) should be very small to estimate the gradient of the expected reward. Learning can therefore not be accelerated by arbitrarily increasing the plasticity factor. In practice, increasing the plasticity factor for one pattern leads to overwriting of previously acquired weight changes induced by other patterns. Not all gradient rules are equally sensitive to this scaling problem. A learning rule that for small *η* would follow the reward gradient in a WTA network, is particularly vulnerable to a large increase of the plasticity factor. With a large *η* the voltage response in the output layer becomes close to an all-or-none structure, and this reduces the difference between the voltage and the stochastically generated WTA spike structure. Since it is this difference that drives plasticity, learning quickly saturates for the presented pattern, and so does it also for the similar other patterns.

As a remedy to the saturation problem for high plasticity factors, the difference term can be enhanced by suppressing the competition term in the voltage accessible to the synapses, without introducing a deteriorating sign change in the plasticity rule. Even though the output spike pattern is established based on the lateral competition, the learning rule ignores this and calculates the difference to the target as the somatic voltage would have been produced only by the feedforward input without lateral inhibition. This leads to a competition-agnostic learning rule that teaches the feedforward afferents to reach the target by them alone – it is a ‘learn to do it yourself’ signal (Fig. 3).

**Fig 3.**
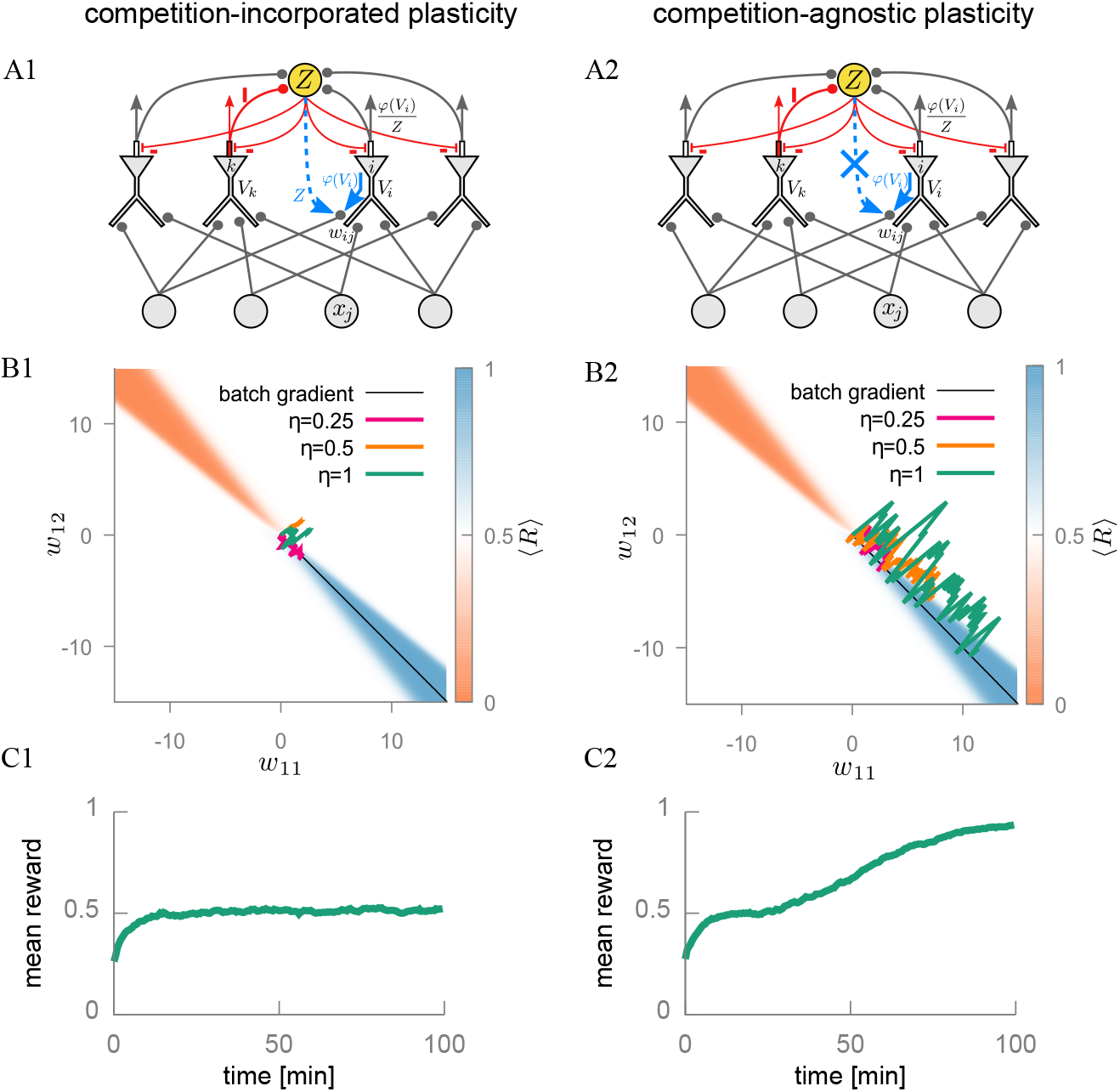
Competition-agnostic plasticity allows for fast learning without saturation. (A) The winner-take-all mechanism is implemented via global inhibition at the axon initial segment, triggered by the first spike of an output neuron (here neuron *k*, red) that inhibits via a global inhibitory neuron (orange) the others from spiking. This leads to the output rate normalized by the total relative firing rate *Z* across all output neurons (Eq. 11) for the competition-incorporated rule (A1, dashed blue arrow) but not in the competition-agnostic rule (A2, crossed dashed blue arrow). (B) Evolution of the weights gets stuck for the competition-incorporated (B1) but not for the competition-agnostic rule (B2). (C) Correspondingly, for high learning rates *η* performance gets stuck for the competition-incorporated (C1), but not for the competition-agnostic rule (C2).

On the implementation level the separation of the lateral inhibition from the forward drive is achieved by deferring the WTA-mechanism with the lateral inhibition to the axon initial segment. The synapses will then predict the spike activity based on the unperturbed local dendritic voltage alone. In fact, because soma and dendrites represent a deep current sink viewed from the axon initial segment, the dendritic voltage is only barely influenced by the voltage in the axonal segment. Yet, the spikes generated there backpropagate to the dendrites where they can be read out by the synapses to identify the output spike rate (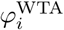, see Eq. 3 and Fig. S1 and Suppl. Mat.). Inhibition of the axon initial segment that shunts the spike trigger mechanism has been found at various types of neurons (Somogyi et al., 1983; Douglas and Martin, 1990).

### Consolidation by selecting informative learning events

High plasticity factors lead to fast learning, but in general also to fast forgetting by overwriting synaptic weights relevant for previous memories (Fusi et al., 2005). To keep the relevant information in the synaptic weight structure, our rule selects informative events for writing over the quickly decaying fast weight components (–*λ_w_w^f^*) into the non-decaying components *w^s^*. This is done by two criteria, one on the strength of the fast weight components themselves, and one on the average postsynaptic voltage (Fig. 4).

**Fig 4.**
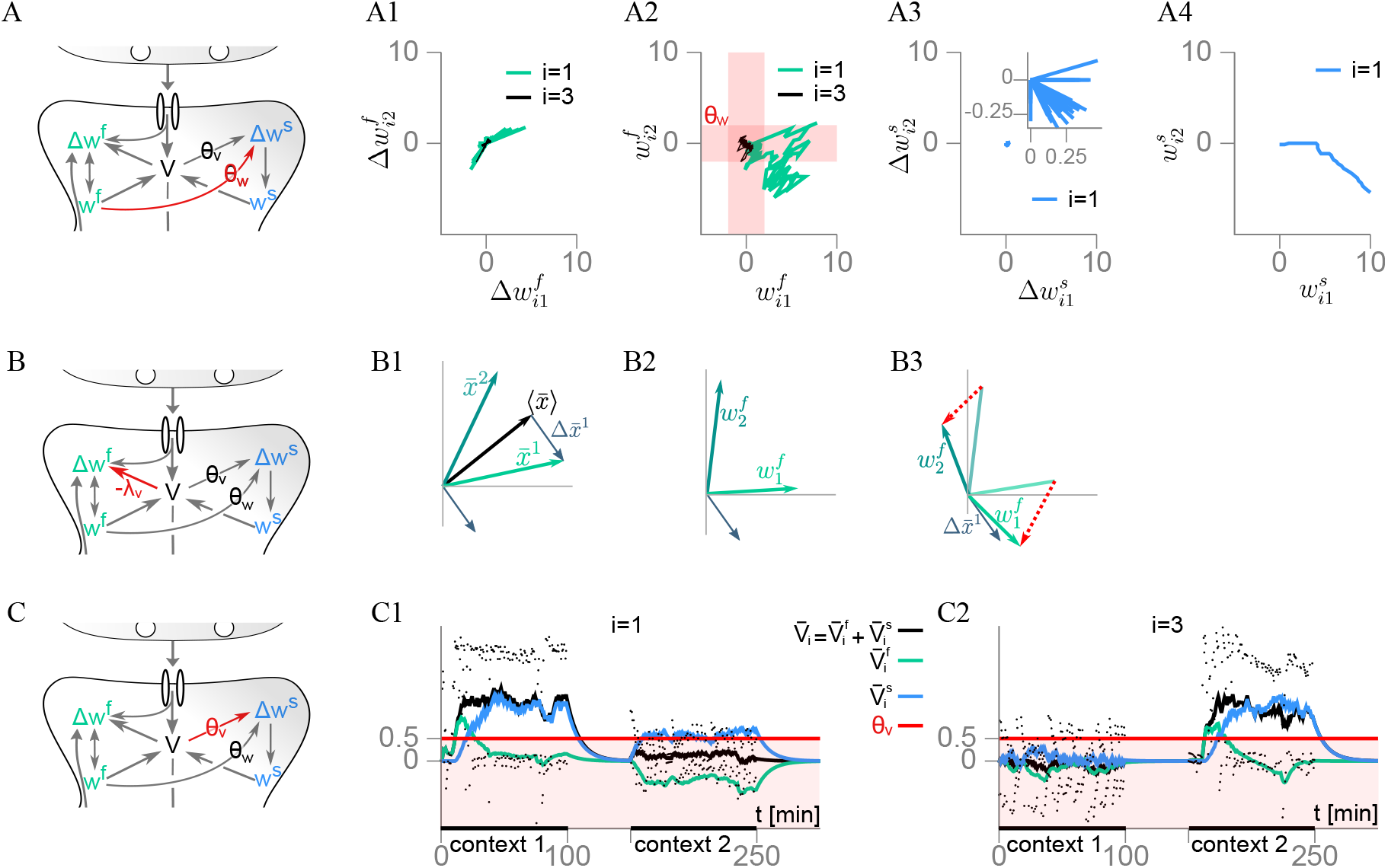
Selecting informative events for consolidation. (A) Only large enough fast weight components (the ‘tagged’ ones) are eligible for consolidation. (A1) The first 100 weight changes 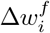 induced by presenting patterns from class 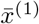 and 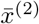, for the two output neurons *i*=1 (green) and *i*=3 (black) and the presynaptic neurons *j* = 1, 2; see also Fig. 1A, B. (A2) The weight changes are summed up, and if a component exceeds the threshold 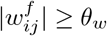 (i.e. if outside the corresponding red shaded stripe), it will be transcribed into the slow component. (A3) The slow weight changes induced by the suprathreshold fast components, 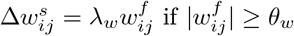 (Inset: zoom-in). (A4) Summing up the slow weight changes 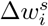 yields an improved gradient estimate 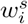 as compared to 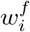 see A2. (B) The voltage penalty minimizes weight overlaps. (B1) The first two input classes 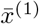 and 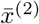 (projected to the first two dimensions) have a large overlap, as expressed by the large mean vector 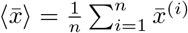 leading to the decomposition 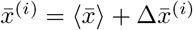 (shown for *i* = 1). (B2) Without voltage penalty term in the energy function (λ_*V*_ = 0), the weight vectors 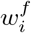 inherit some overlap from the input patterns. (B3) By penalizing systematic voltage deflections in one direction (*λ_V_* > 0), the weight vectors become roughly orthogonal to the mean pattern and instead align with the deviations form the mean, 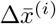. (C) Only events with strong enough mean depolarization (producing ‘plasticity related proteins’) are informative about the context and eligible for consolidation. (C1) The low-pass filtered voltage of neuron 1 (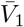, black) exceeds the consolidation threshold (red) during context 1 as it should respond to input class 1, but it remains sub-threshold (red shaded region) during context 2 and hence the new context does not touch the previously consolidated weights of neuron 1. This would be different if the condition for consolidation were imposed on the non-averaged voltage (*V*_1_, black dots), or if the voltage were driven by the slow weight component alone (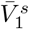, blue). Yet, because neuron 1 is never the correct winner, the fast component quickly learns that neuron 1 should be inactive in context 2 (averaged voltage induced by this fast component, 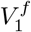, green, is negative) and its consolidation is prevented. (C2) Same as in C1 but for neuron 3 that undergoes weight consolidation only in context 2.

To motivate the first selection criterion, we note that changes of the fast component, summed across the pattern presentations in time, 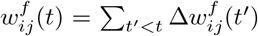, yield a more robust gradient estimate than individual updates 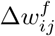 themselves. One idea to improve the estimate is therefore to consider the fast weight components as samples of the gradient, and update the slow component not by 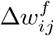 but by the potentially better gradient estimates 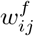. These summed estimates are only better if they occurred often enough in the same direction, and this can be expressed by the criterion 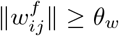. This justifies the criterion for the slow weight update (cf. Eq. 4)

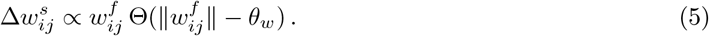

To motivate the second selection criterion we consider the penalty term in the utility function that punishes strong voltage deflections on |*V_i_*|, see Eq. 1. At the level of the gradient this leads to the forgetting term in the weight update of the form, 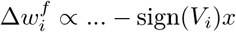, where 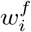 is the weight vector targeting the postsynaptic neuron *i*. It subtracts away common components in the inputs *x* from 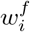 until, on average, this term vanishes, i.e. 〈sign(*V_i_*)〉 ≈ 0, where 〈·〉 denotes the average over a long sequence of pattern presentations. As a consequence, 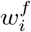 becomes roughly orthogonal to the average input pattern, 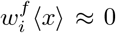, making the voltage 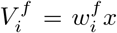 informative about pattern differences (Fig. 4B).

Imposing a voltage threshold for the transcription into the slow component selects the appropriate neuron for which synapses are consolidated. Applying the threshold condition on the temporal mean instead of the instantaneous voltage, favors the consolidation of associations that are encountered in the temporal vicinity of strong and lasting depolarizations. This becomes important when contexts are switched and patterns that were similar in the previous context now cause a strong but erroneous depolarizations that should not be consolidated (Fig. 4C, black dots). Fortunately, due to the fast learning in *w^f^*, an incorrect strong depolarization will only arise a few times in a row, and by imposing the threshold onto a consistently high depolarization, only the correctly activated neurons become selected. This justifies the voltage-based consolidation criterion,

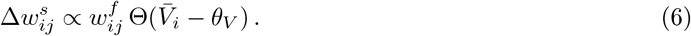

### Synaptic tagging, capture and consolidation

We next interpret the consolidation mechanisms in terms of biological quantities arising in the synaptic tagging & capture framework (Frey and Morris, 1997; Redondo and Morris, 2011). We postulate that whenever the absolute value of the fast weight component crosses a threshold, |*w^f^*| ≥ *θ_w_*, a synaptic tag in that synapse is set that remains active as long as |*w^f^*| is above threshold. Next, if the low-pass filtered postsynaptic voltage is above a threshold, 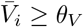, the translation of plasticity-related proteins (PRPs) is triggered that move up the dendritic tree. A synapse with an active tag captures PRPs and a fraction of the fast weight component *w^f^* is transcribed into the slow component (Fig. 5A).

**Fig 5.**
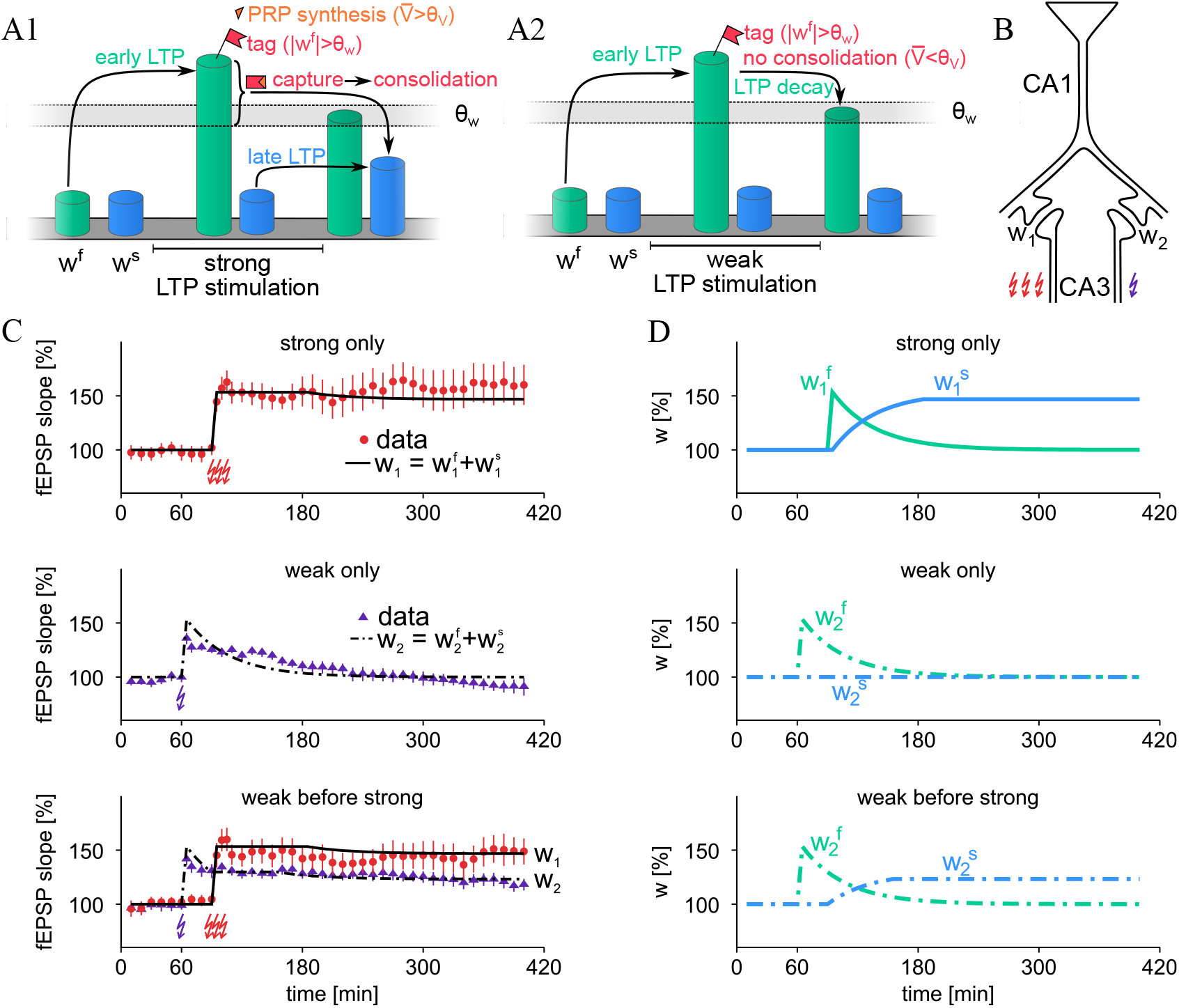
Model accounts for synaptic tagging *in vivo*. (A1) Early LTP in the fast weight component *w^f^*, e.g. elicited by weak or strong tetanus, sets a local synaptic tag if *w^f^* crosses a threshold, |*w^f^*| ≥ *θ_w_*. Strong and lasting postsynaptic depolarization triggers the synthesis of plasticity-related proteins (PRPs), 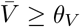. When a tag captures available PRPs (red & orange symbols), the fast weight component is consolidated in the slow component *w^s^* (late LTP, Eq. 4). (A2) When no PRPs are available, the early LTP decays without consolidation (Eq. 3). (B) Experimental setup from Shires et al. (2012). Two afferent pathways projecting to a common CA1 pyramidal neuron (modeled with synaptic strengths *w*_1_ and *w*_2_) are stimulated by a strong tetanus (3 red flashes, such that 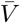 crosses threshold) and weak tetanus (1 purple flash, without *θ_V_* crossing but still triggering postsynaptic spikes). (C) Top: Strong tetanization of one pathway only results in long-lasting changes in the data and the model. Middle: For weak tetanization of a single pathway, the synaptic strength (that is the sum of the fast and slow component) decays back to baseline. Bottom: If the weak tetanization of pathway 2 (purple) is followed by a strong tetanization of pathway 1 thirty minutes later (red), the decay of the synaptic strengths in pathway 2 is stopped. (D) Evolution of the separately shown fast and slow weight components of the strongly (solid, *w*_1_) and weakly (dashed, *w*_2_) stimulated pathways.

Based on this interpretation, we can reproduce results from synaptic tagging experiments in vivo, where electrodes were placed bilaterally in CA3 to stimulate independent synaptic inputs targeting a common population of postsynaptic neurons in CA1 (Fig. 5B, Shires et al. (2012)). A strong tetanus protocol applied on pathway 1 (such that the postsynaptic voltage 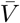 crosses threshold and strong postsynaptic firing is elicited, Methods) directly consolidates the early LTP in the that same pathway, provided the synaptic tag was activated (i.e. 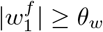, Fig. 5C, top). If pathway 2 is stimulated weakly (such that 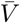 does not cross *θ_V_*, but postsynaptic spikes still triggered) the induced early LTP decays back to baseline again (Fig. 5C, middle). However, if within 30 minutes after (or before) the weak stimulation the second pathway is strongly stimulated such that now 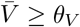 (and PRPs are expressed), also the first pathway gets consolidated, provided there the tag is activated by the previous (or following) weak stimulation (Fig. 5C, bottom).

The synaptic tagging and consolidation experiments are astonishingly symmetric with respect to LTP and LTD (Sajikumar and Frey (2003, 2004), see Fig. 6). This may be caused by the fact that in all experiments plasticity was induced by presynaptic stimulations of excitatory afferents, although for LTD the overall stimulation was weaker. Whether LTP or LTD is induced may depend on the elicited postsynaptic spike rate. In our model, the sign of the plasticity induction depends on whether the somatic spiking activity is higher or lower than the dendritic prediction (yielding a positive or negative sign in the error term 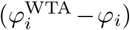 entering in the update 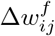, see Eq. 3). To describe the unsupervised character of the experiments, the reward prediction error 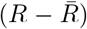 was set to a constant value (= 1). To reproduce the additional experimental data we assumed that, for simplicity, no postsynaptic spike activity was triggered during the LTD stimulations (in contrast to the LTP stimulations), while both the strong LTD and LTP stimulations generated a local low-pass filtered dendritic voltage that was suprathreshold (in contrast to weak LTD and LTP – weak LTP stimulations may trigger more global depolarization that still causes postsynaptic spiking).

**Fig 6.**
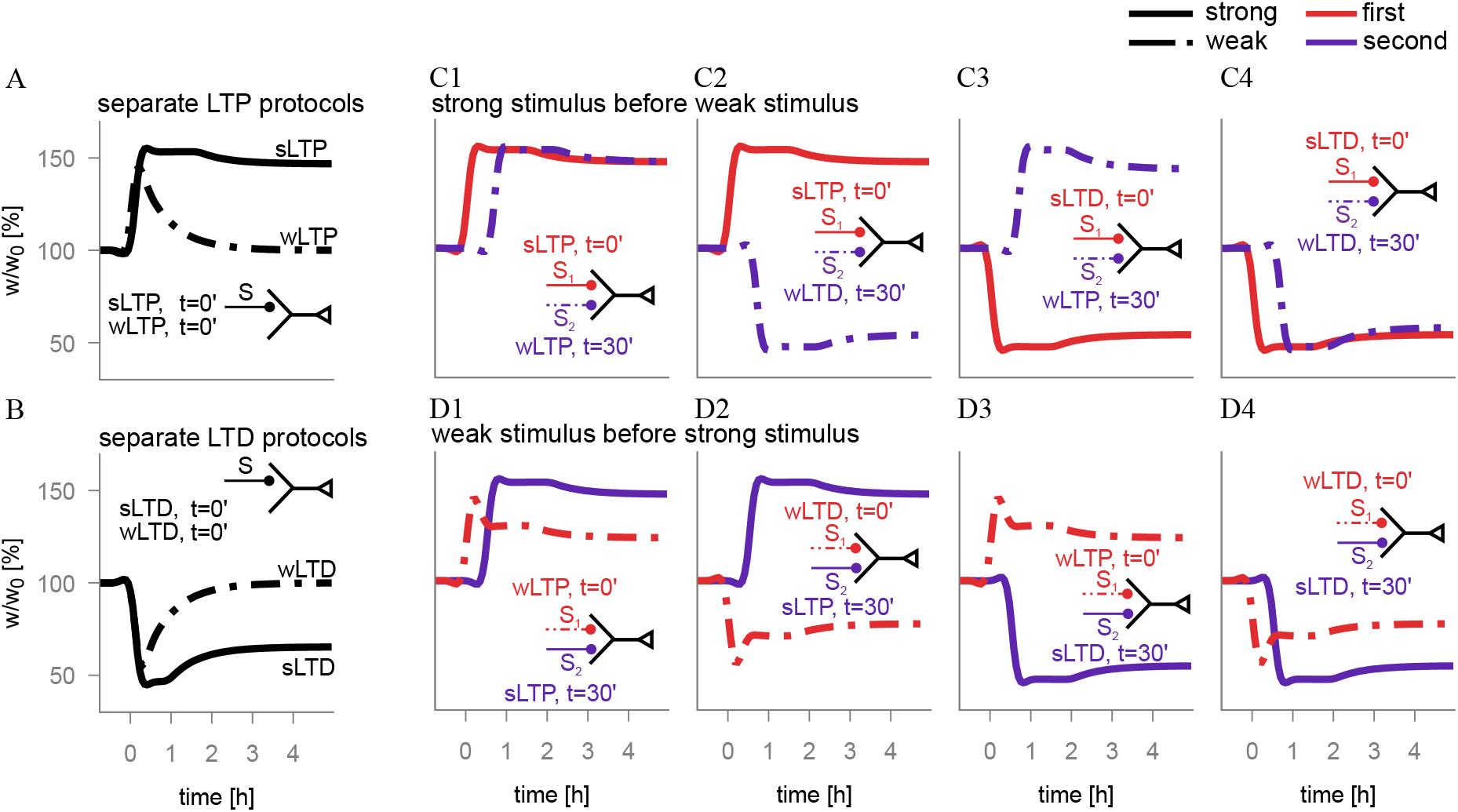
The model also accounts for LTD tagging and cross-tagging. For the corresponding experimental data see below. (A) Strong (solid) and weak (dashed) LTP stimulation applied in separate ‘experiments’ causes a strong lasting and a weaker decaying weight change, respectively (shown is *w*= *w^f^* + *w^s^*). (B) Same as in A, but for LTD protocols. (C) Strong stimulation applied 30 minutes before weak stimulation on different pathways rescues the weight decay after weakly induced early LTP and LTD. (D) Same as in C, but for the strong stimulation applied 30 minutes after the weak plasticity induction. Insets summarize the stimulation protocols that have also been applied in the experiments. A: Frey and Morris (1997), B: Sajikumar and Frey (2003), C1: Frey and Morris (1997), C2-C4: Sajikumar and Frey (2004), D1: Frey and Morris (1998), D2-D4: Sajikumar and Frey (2004)

Figure 6 summarizes the various experiments that were reproduced by our plasticity model of Eqs. 3 and 4. These experiments also include the so-called cross-tagging according to which a strong LTP protocol consolidates weakly induced LTD, and a strong LTD protocol consolidates a weakly induced LTP, be the strong protocol applied before or after the weak induction (Fig. 6C, D).

### Fast learning deteriorates pattern discriminability when mixing easy and difficult tasks

So far we have shown how fast learning in different contexts with similar inputs is possible while reducing context interference. However, fast learning comes with a price. If in a given context classification tasks of unequal difficulties are mixed, learning slows down. This is indeed a well known phenomenon in psychophysics, called roving (Tartaglia et al., 2009b; Parkosadze et al., 2008) or interleaved learning (Flesch et al., 2018).

To exemplify these phenomena, we consider a perceptual bisection task with two different stimulus types characterized by offsets (*ϵ*) from the middle line chosen around larger (easy) or smaller (difficult) means (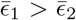, Fig. 7A). After each stimulus presentation, participants have to tell in which direction they perceived the offset. To model this task we set up a network with 100 input neurons and two output neurons (Fig 7B) with an all-to-all connectivity from input to output neurons. The firing rates of either the first or second half of the input neurons are sampled around the mean 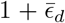, while the other half is sampled around 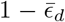 with *d* = 1 or 2. The mean offset 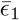 or 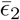 codes for the easy and difficult stimulus type, respectively. If the first half of the input neurons fire on average with a higher firing rate, the first output neuron needs to be activated to get reward (*R*= 1), ifthe second halfofthe neurons fire on average more, the second output neuron needs to be activated. If incorrectly classified, reward is omitted (*R* = 0).

**Fig 7.**
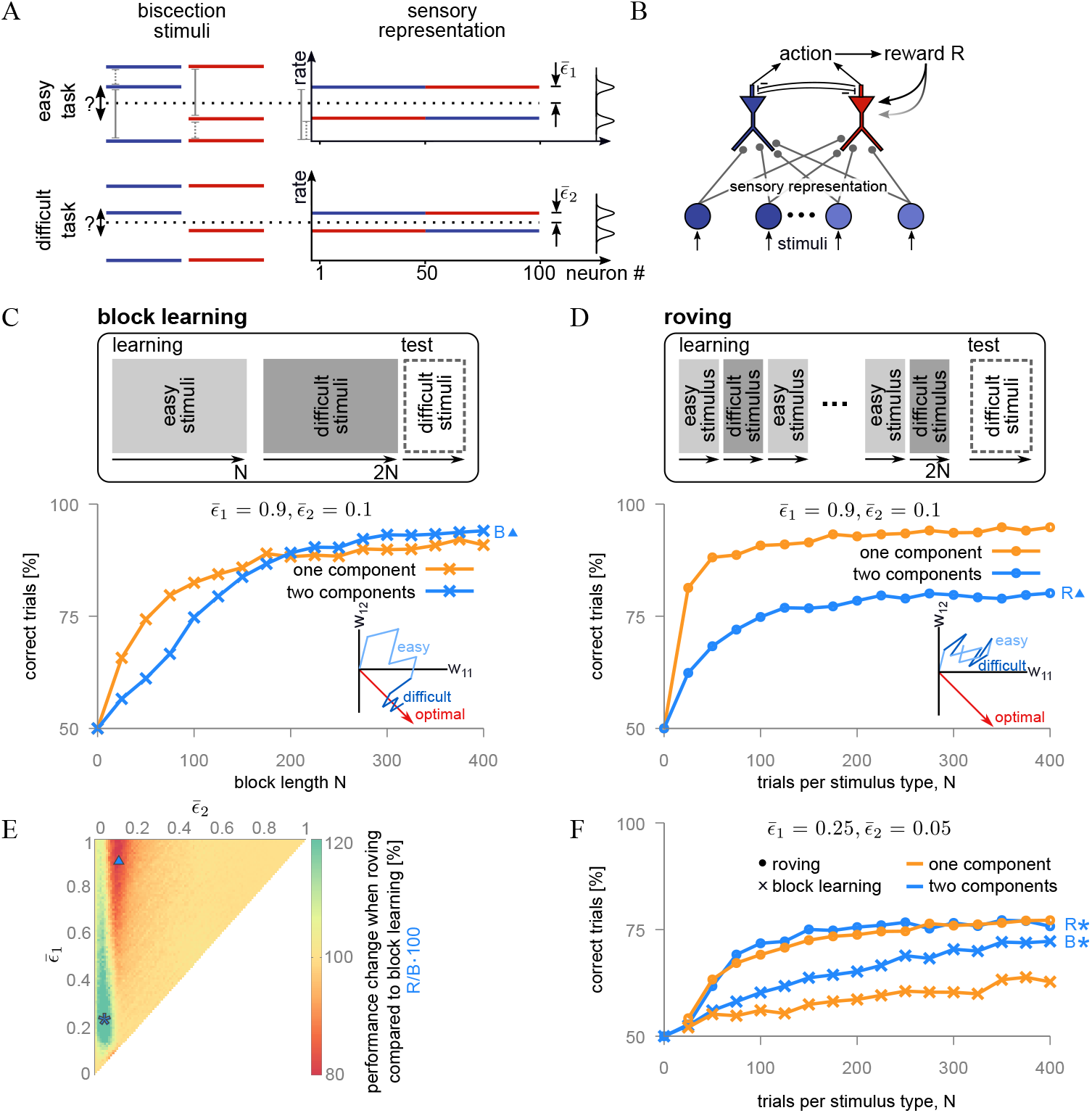
2-component rule accounts for roving. (A) In a bisection task participants have to decide whether the midline is offset to the top or bottom (left). In the model (right), either the first (1-50, blue) or second (51-100, red) half of the sensory neurons have a higher firing rates. Two stimulus types are considered, an easy one with a large midline offset (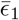, right top) and a difficult one with a small offset (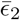, right bottom). (B) The model network has 100 sensory neurons and two action neurons (coding for the up-or down shift from the midline) that mutually inhibit each other with their ‘first spike’. If the first (second) half of the input neurons have a higher mean firing rate, the first (second) output neuron has to be active to get global reward *R* = 1, otherwise *R* = 0. (C) If the easy and difficult stimulus types are learned block-wise (*N* trials per block), both the 1- and 2-component rules learn the task. Performance is shown only for the difficult stimulus type. Inset: Example of a 2-component weight trajectory. During the first block of easy stimuli (light blue) the weight jumps strongly around the optimal weight (red), and converges during the block with the difficult stimuli (dark blue, *N* = 7). (D) Easy and difficult bisection stimuli are presented in random order (roving). Compared to block learning, performance for the difficult stimulus type improves more strongly than with the 1-component rule and drops with the 2-component rule, in accordance with the psychophysical experiments. Inset: The large fluctuations caused by weight updates in response to the more varying easy stimuli (light blue) cannot be corrected by the interleaved difficult stimuli (dark blue) and the fluctuations around the optimal weight (red) remain larger as compared to the block-wise learning in C. (E) Performance change for roving compared to block-learning, for any pair of task difficulties 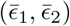, using the 2-component rule. (F) If the difficulty of both types is increased while becoming more similar, both the 1- and 2-component rules predict that roved learning outperforms block learning.

If the easy stimuli are presented in a first block, followed by the difficult stimuli in a second block, learning is possible for both the 1- and 2-component rules (Fig. 7C, similar results for reversed block order, not shown). The same learning rates were used as the ones optimized for the context learning task (cf. Fig. 1). If the easy and difficult stimuli are presented randomly interleaved, the 2-component rule performs less well, unlike the 1-component rule (Fig. 7D), analogously to human behaviour (Parkosadze et al., 2008; Clarke et al., 2014). The performance decrease with roving (mixing) occurs because weight changes, in response to an easy stimulus, point further away from the optimal weight due to the larger variance of presynaptic rates around the mean for these easy stimuli. When presented in a block, the large deviations are corrected during the learning of the difficult stimuli that themselves show lower variance (inset of Fig. 7C). If the easy and difficult stimuli are mixed, however, the large variances remain throughout learning and prevent a convergence towards the optimal weight (inset of Fig. 7D). Weight updates also become smaller during learning because the modulating reward prediction error typically becomes small (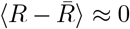 as *R* = 1 most of the time). Learning in the roving condition with the 1-component rule is better because the learning rate optimized for the previously studied context learning is smaller, paying out for roving, but also leading to catastrophic forgetting.

Interferences between stimulus types diminish when the two stimulus types become more similar in difficulty. Since reinforcement learning follows the principle of exploration and exploitation, a certain level of exploration in the weight space, for instance caused by mixing of different stimulus types, may be beneficial. This is in fact what occurs when the difficulties for the two stimulus types become similar and they are randomly mixed to boost stochastic weight fluctuations (Fig. 7E, star). Consequently, the high variance weight updates from the easier stimuli, that would hinder discrimination for the difficult stimuli, now rather improve the performance in the roving scenario for the 2-component rule as compared to block learning (Fig. 7F, blue).

## Discussion

We demonstrated that catastrophic forgetting in continual learning tasks can be prevented by using two synaptic weight components, a fast and a slow one. The dynamics of the fast component are derived from a stochastic gradient procedure on a reward-based utility function. The utility incorporates the expected reward and a sparsity constraint on both, the postsynaptic voltage and the fast component. Since the change of the fast component follows the utility gradient, this weight component maximizes the expected reward under the constraint of small weight modifications and small voltage deflections. The sparsity constraint on the voltage implies an orthogonalization of the pattern representation, and the sparsity constraint on the weight implies a passive decay of the fast component. Both the orthogonalization and the decay help to reduce interference when new associations for similar stimuli have to be learned. To not forget the old associations, part of the decaying fast weight component is consolidated in a slow weight component. An additional selection process on the strength of the fast weight component and the amplitude of the voltage deflections insures that only the ‘informative’ events are memorized. This selection process nonlinearly amplifies the two sparsity constraints on the weight and the voltage: only when the fast weight component exceeds a threshold, and only when the averaged postsynaptic voltage is simultaneously above a voltage threshold, the fast component is consolidated.

We show that the 2-component gradient rule, unlike 1-component rule(s), prevents catastrophic forgetting across subsequent contexts in which classification tasks with similar input patterns but different outputs have to be learned. Intriguingly, after a pause (of roughly an hour biological time) the test performance of the model network is even better than before the pause. This is explained by the decay of the fast component that leads to a forgetting of the sample-specific noise, while the class-relevant information is consolidated akin to the consolidation and the semantization of memory during sleep (Stickgold, 2005). We also show that the consolidation criteria on the strong weight component and the large voltage deflection can be interpreted in terms of synaptic tagging and capture (Frey and Morris, 1997; Redondo and Morris, 2011). The synaptic tag & capture hypothesis posits that a weak synaptic stimulation that triggers a tag during the induction of early LTP/LTD – in our model the threshold-crossing by potentiating/depressing the fast component – is captured by a strong stimulation within some minutes before or after – in our model the threshold-crossing of the low-pass filtered postsynaptic voltage causing consolidation. Notably, our model implicates that reward and consolidation arise also from weak LTP to strong LTD and from weak LTD to strong LTP, just in the same way as in the real experiments dopamine is implicated and cross-tagging between potentiation and depression is observed.

The strategy of consolidating only large weight components associated with high voltages leads to an overweighting of strong stimuli. A similar overweighting of large magnitudes is also observed when humans solve a categorical decision task under limited cognitive resources, e.g., induced by time constraints (Spitzer et al., 2017). In economics, the selective integration that discards low-value options is known as economic irrationality (Tsetsos et al., 2016), reminiscent of discarding low-activity events when learning associations across time. In a similar way as economic irrationality remains optimal under constraints, our learning rule remains hill-climbing on the utility function in the competition-agnostic version that discards winner-take-all information among the output neurons. Upon reward, the competition-agnostic rule considers the postsynaptic activity as a target, irrespectively of how the activity is produced, and by this virtue prevents an early saturation of learning even for very high learning rates. Turning reinforcement learning into a target-based learning appears as a trick to speed up learning. The ultra-fast learning rate for the competition-agnostic learning rule is only possible with the 2-components that enable the selective decay of the ‘non-informative’ weight changes while consolidating the ‘informative’ ones.

Fast learning with selective consolidation comes with a price. Fast learning is enabled by a transient synaptic memory buffer that has an intrinsic time constant. Its downside is exposed when intermixing tasks of unequal difficulties that fill up the buffer with unspecific information. Randomly mixing easy and difficult stimuli (‘roving’) in a classification task hampers learning with the 2-component rule, but not with the rate-optimized 1-component rule. A similar phenomenon is observed in perceptual discrimination tasks when bisection stimuli of unequal difficulties are randomly intermixed (Otto et al., 2006; Parkosadze et al., 2008; Clarke et al., 2014). In our model performance degrades because the easy stimuli simultaneously trigger suprathreshold weight changes and suprathreshold voltages that are then prematurely consolidated, even when these changes are not accurate enough to correctly classify the difficult stimuli. A previous model was explaining the roving phenomenon by an imprecise critic in the context of reinforcement learning (Herzog et al., 2012). When mixing tasks of different difficulties, the critic induces a synaptic drift towards either LTD for the difficult and LTP for the easy task, or vice verse. In our model, learning is hampered not due to a systematic drift, but due to the early consolidation of erratic weight changes triggered by the easy task.

The model we presented can be seen as one of many attempts to fight catastrophic forgetting during continual learning with sophisticated synaptic consolidation mechanisms. The synaptic cascade model (Fusi et al., 2005; Benna and Fusi, 2016; Kaplanis et al., 2018) postulates hidden states that, different from our model, do not contribute to the effective synaptic strength. Others, more globally defined learning rules have been suggested that keep synaptic parameter changes small if the same parameters were previously engaged in a performance increase (Kirkpatrick et al., 2017; Zenke et al., 2017). A yet more global criterion for learning rate adaptation has been suggested that prevents catastrophic interference using conceptors that seek to orthogonalize the pattern representation in different contexts (He and Jaeger, 2018).

Our gradient-based 2-component plasticity model yields a synaptic explanation of well-known learning strategies. (1) Context switches should not be faster than the decay time of early LTP/LTD (some ten minutes) to prevent interferences induced by the not-yet-decayed fast weight components. (2) A learning break without stimulus exposures helps to semanticize memories through forgetting of non-systematic, random stimuli that did neither trigger suprathreshold fast component changes nor suprathreshold neuronal activities. (3) Mixing tasks of unequal difficulties may block the learning of the difficult task by premature weight consolidations triggered during the easy task. On the synaptic level we predict that consolidation thresholds are dynamically adapted to the statistics of the fast weight- and postsynaptic voltage-distributions, preventing catastrophic forgetting in a volatile environment, and endowing individual synapses in deep networks with a local consolidation mechanism.

## Methods

Throughout the paper a single-layer network with an all-to-all connectivity between input and output layer is used. Each postsynaptic neuron receives from each presynaptic neuron input *x_j_*(*t*) through synapses with weight *w_ij_*(*t*). The postsynaptic voltage *V_i_*(*t*) can elicit a spike which mediated by strong lateral inhibition suppresses all other postsynaptic neurons from spiking upon onset of a new input. The probability that a spike is observed in neuron *k* is given by the normalized Poisson firing rate,

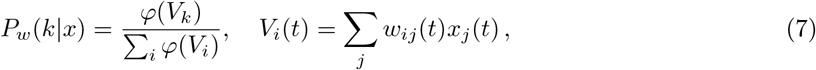

where *φ* denotesthelogisticfunction, *φ*(*x*) = *φ*_max_(1+exp(–(*x*–*α*)/*β*))^-1^. The strong lateral inhibition generates a WTA dynamics. As a result one gets for each presented input pattern an output vector of the form *y^k^*(*t*) = (0, …, 0, 1, 0, …, 0) where 1 is at position *k*. The environment is evaluating the output and issues a global binary reward signal *R* to the network. For each input class there is exactly one output which results in *R* = 1, all other responses give *R* = 0.

### The competition-incorporated rule is gradient ascent

Using the 2-component rule, each synaptic weight consists of two components, a fast component 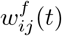 and a slow component 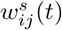. The total synaptic weight *w_ij_*(*t*) is the sum of the fast and the slow component, 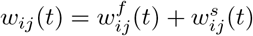.

The update of the fast component, Eq. 3, is derived from the gradient ascent rule of the utility function in Eq. 1. The expected reward is given by

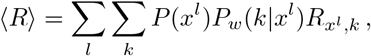

where the sums run over all input pattern indices *l* and potential winning units *k. R*_*x*^*l*^,*k*_ denotes the value of reward given for input-response pair (*x^l^, k*). The gradient of 〈*R*⌣ with respect to *w_ij_* can be calculated by using the log trick,

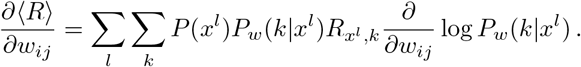

The derivative term can be computed with *P_w_*(*k*|*x^l^*) from Eq. 7 as

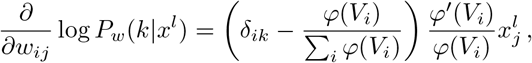

where *k* being the winning unit and the Kronecker delta *δ_ij_* is 1 if *i* = *k* and 0 otherwise. To minimize the variance of the reward estimate we center the effective reward around its approximate mean 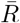,

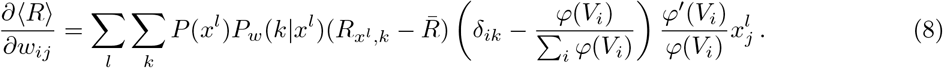

The terms involving 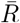 do not originate from the derivative but since they add to 0 they can be added for convenience. If (*x^l^, k*)-pairs are sampled with probability *P*(*x^l^*)*P_w_*(*k*|*x^l^*), the learning rule based on gradient ascent of *U* is given by

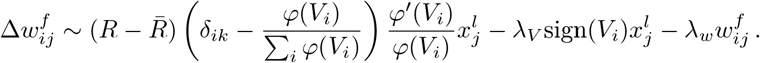

We next assume that the neuronal transfer function *φ*(*V_i_*) saturates for large arguments at *φ*_max_. By setting 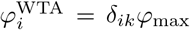 (i.e. 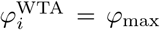 if *i* is the winner, *i* = *k*, and 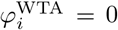 for *i* ≠ *k*), 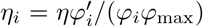 for some global learning rate *η*, and *Z* = Σ_*i*_*φ_i_*/*φ*_max_, we end up with the learning rule

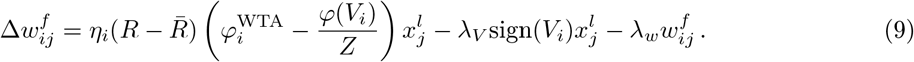

Time-averaged quantities as 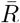 and 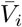 are calculated according to 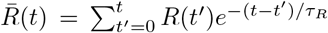, and analogously for 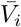. We set *τ_R_* = *τ_V_* = 10 trials, with a trial duration of 30 s. The results do not qualitatively change when choosing half or twice as large time constants. The fast and slow weight components are updated every 30 s biological time (including the ‘breaks’ when *R* = 0 and *x* = 0 by definition).

### The competition-agnostic rule is hill-climbing

The reward-based component of the competition-agnostic learning rule,

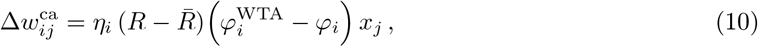

is stochastic hill-climbing on the expected reward 〈*R*〉. This is because the reward-based component of the competition-incorporated learning rule

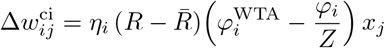

according to Eq. 8 is stochastic gradient ascent on 〈*R*〉, and the two updates are always within 90°, Δ*w^ca^* Δ*w^ci^* > 0. The latter scalar product is positive because for both rules the winner is the same, and 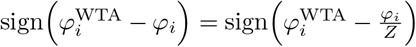 for all *i*. Hence, the averaged update vectors are still within 90^°^, while the averaged competition-incorporated update points in the direction of the reward gradient, 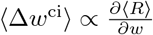. Adding the two vectors Δ*w^ca^* and Δ*w^ci^* to the same penalty gradient 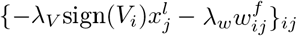 does only decrease the angle. We conclude that the competition-agnostic rule is hill-climbing on the utility function *U*.

### One-component rule

For comparison, all simulations were repeated with a 1-component rule that consisted of the fast component *w* = *w^f^* only (Eq. 3), but with *λ_w_* = 0 and hence *w^s^* = 0. Parameters of the 1-component rule were optimized independently from the 2-component rule.

### Associative task

In the associative task, we consider input classes 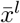 with each class requiring its desired output defined by the exclusive activity of a specific output neuron. Input patterns, which are defined as noisy samples *x^l^* around 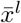, are presented each after the other. The goal is to change the synaptic weights such that for each input class the desired output is obtained. For a pattern of the class *l* the instantaneous Poisson firing rate of the presynaptic neuron *j* is

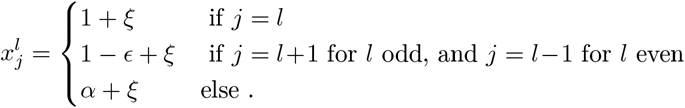

where *ξ* is independent Gaussian noise with zero mean. Two input classes form together a context. Contexts are learned in succession without repetition. Within a context, the input patterns of the two classes are presented repeatedly and in random order. The number of presynaptic neurons in the network corresponds to the number of classes, the number of postsynaptic neurons is fixed at 10. Reward is given if for input pattern *x^l^* output neuron *l* is exclusively active (*R* = 1). Otherwise the reward signal is omitted (*R* = 0). After learning a context, a phase with a null input but unchanged dynamics was on. Performance of learning is evaluated during a test period at the end of the task in which all input patterns are presented again though now without synaptic plasticity. Performance is measured by the percentage of correct trials during the test period. Parameters were set to *η* = 1, λ_*w*_ = 0.012, λ_*V*_ = 0.45, *θ*_*w*_ = 2, *θ*_*V*_ = 2.

### Tagging experiments

In tagging experiments a network with five presynaptic neurons and one postsynaptic neuron was used. Only one of the presynaptic neurons was stimulated at a time. Simulation were run with four stimulation protocols: strong LTP, weak LTP, strong LTD and weak LTD. Stimulation strength was regulated by the input duration. Strong LTP and LTD lasted 15 minutes biological time and weak LTP and LTD lasted 5 minutes biological time. The stimulation amplitude was kept fixed for all four protocols.

For LTP stimulations the teacher term *φ*^WTA^ in Eq. 3 was constantly set to *φ*max whereas it was set to 0 for LTD stimulations. In all tagging experiments, 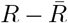 was set to 1. If two protocols were paired, the second stimulation set in 30 minutes after the first stimulation on a second presynaptic neuron.

In experiments, late-LTP was induced in vivo by a strong tetanization consisted of three trains of 50 pulses at 250 Hz, with a 5-min intertrain interval; weaker induction of LTP was investigated by a weak tetanus consisting of 1 train of 50 pulses at 100Hz (Shires et al., 2012). For LTD, see Sajikumar and Frey (2003): Late-LTD was induced using a low-frequency stimulus protocol (LFS) of 900 small bursts (one burst consisted of three stimuli at 20 Hz, interburst interval 1 s, i.e. f=1 Hz, stimulus duration 0.2 ms per half-wave, a total of 2700 stimuli). This stimulation pattern produced a stable long-term depression in vitro for at least 8 h. In experiments in which a weaker induction of LTD was investigated, a transient early-LTD was induced using LFS consisting of 900 pulses (1 Hz, impulse duration 0.2 ms per half-wave, a total of 900 stimuli).

### Roving

To explore the phenomenon of roving a bisection task with two stimulus types were used, an easy and a difficult type. The bisection task consisted of two input patterns which needed to be discriminated. The stimulus types only differed in the difficulty to discriminate these input patterns. For each input pattern, half of the population had mean Poisson firing rate of 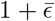, the other half of 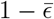, where 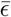 is a stimulus type specific parameter. Each input was modulated by an additive Gaussian noise ξ,

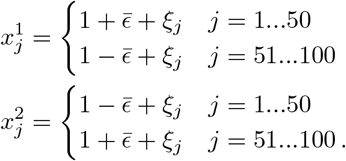

The network consisted of 100 presynaptic neurons and two postsynaptic neurons representing the two possible actions in the experiment of pressing left or right. Reward was given if neuron 1 was activated and the first half of the input population had a higher activity or if neuron 2 was activated and the second half of the population had a higher activity. Two learning scenarios were considered: block learning and roving. In the block learning scenario, easy input patterns were first trained followed by the training of the difficult patterns. During roving, patterns of the easy and difficult type were alternated presented in random order. The total number of pattern presentations was for both learning scenarios equal. In the test period, which followed a phase with null inputs, the percentage of correct trials using patterns of the difficult type was evaluated.

### Simulation details

Simulations were run on the HPC cluster of the University of Bern. For the associative tasks (Fig. 2), all parameters were separately optimized for the 1- and 2-component rule with a multivariate bisection method. The objective function was the sum of the expected reward after 200 simulation runs. Each optimization run was performed with new random values of *α* and *ϵ* and a random number of contexts between two an six. Keeping parameters fixed, simulations were done for combinations with *α* ∈ [0,1] and *ϵ* ∈ [0,1] using step size 0.01. The percentages of correct trials are average values over 200 simulations.

For the tagging experiments (Fig. 5 and 6), parameters were adapted to best fit the in vivo results by Shires et al. (2012) reproduced in Figure 5. For additional protocols and pairings shown in Figure 6 the parameters were kept unchanged.

In the roving scenario (Fig. 7), the same parameter were used as for the associative task. In panels 7C-E each point represents a separate simulation set. Shown values are averages over 200 simulations.

## Acknowledgments

This work was supported by the Swiss National Science Foundation (SNSF, Sinergia grant CRSII2-147636 to MH and WS), and by the European Union’s Horizon 2020 Framework Programme for Research and Innovation under the Specific Grant Agreements No. 785907 (Human Brain Project, SGA2).

## Supplementary Material

### Competition-incorporated plasticity quickly saturates

To formally capture the described problem of large plasticity factors we consider the exact gradient of the expected reward that includes a global normalization *Z* (Methods),

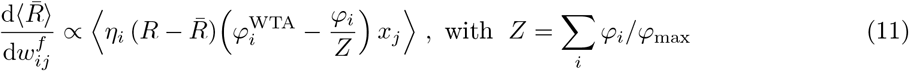

for *φ_i_* = *φ*(*V_i_*). As compared to the approximate stochastic gradient (Eq. 3), the normalization by *Z* appears. This normalization makes the ratio *φ_i_*/*Z* identical to *φ*_max_ times the probability that neuron *i* is the winner (with *k* being the index of the winner neuron and 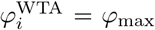 for *i* = *k* and 0 else). If for a synaptic update a sample of the right-hand side of Eq. 11 is used with a high plasticity factor, 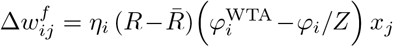, the winner probability quickly becomes all-or-none, 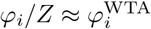, and learning will saturate. Leaving out the normalization by *Z*, however, keeps Δ*w_ij_* away from 0, without changing the sign of Δ*w_ij_*. Overall, this yields a plasticity rule (Eq. 3) that allows for speeding up the learning by increasing *η_i_*. On average, the update still stays within 90° of the true gradient as the synaptic bias does not invert the sign of the weight change, and thus the competition-agnostic rule remains hill-ascending on the utility function (Methods, Fig. S1).

**Fig S1.**
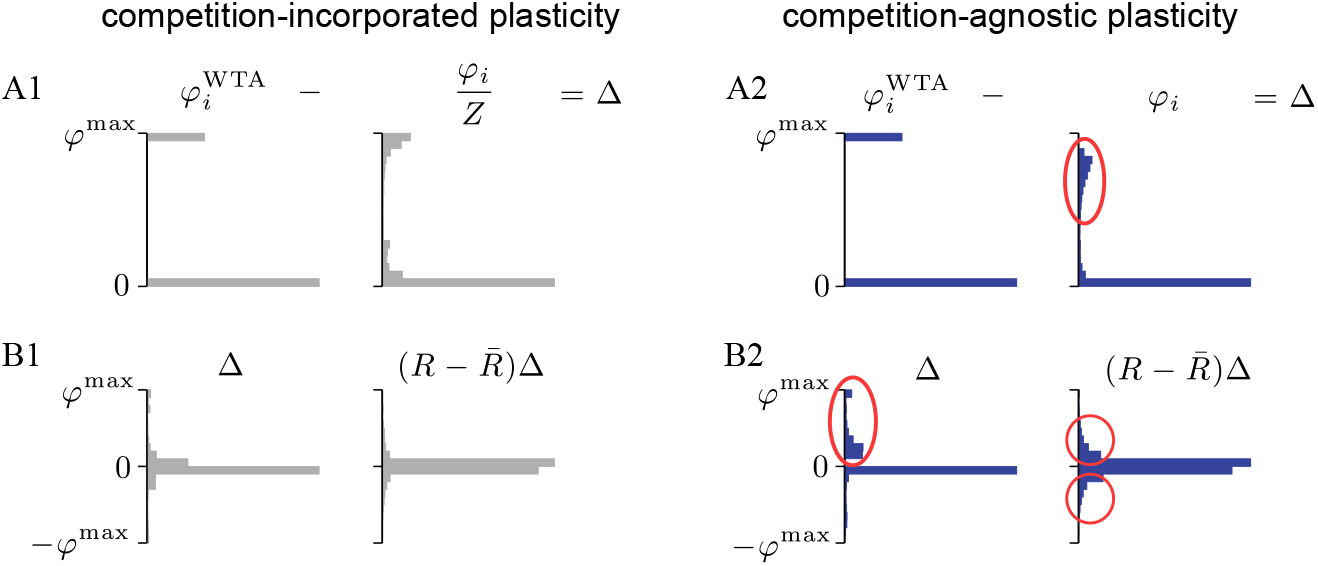
Competition-agnostic plasticity create non-vanishing learning signals. (A1, B1) Learning stalls for high *η* since the error in the competition-incorporated rule, 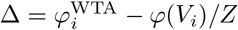, mostly vanishes. This arises because for fast learning the output quickly converges to a one-hot representation, such that *φ*(*V_i_*)/*Z* gets close to 1 (for *i*=*k*) or 0 (for *i*≠*k*), as this is by construction the case for 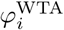. (A2, B2) This is different for the error term 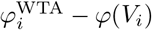 of the competition-agnostic rule where *φ*(*V_i_*) is not normalized (A2, oval), leading to non-vanishing errors Δ (B2 left, oval). Although the error term Δ is now transiently unbalanced, the product 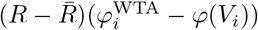 across synapses becomes balanced again (B2 right, two circles) because 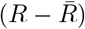 is balanced, preventing a deteriorating weight drift. In fact, the competition-agnostic rule remains hill-climbing (Methods).

